# Massively Parallel Dissection of RNA in RNA-protein interactions *in vivo*

**DOI:** 10.1101/2022.06.13.495920

**Authors:** Evan P. Hass, Yu Hsuan Lee, Will Campodonico, Yong Kyu Lee, Erika Lasda, Jaynish S Shah, John L. Rinn, Taeyoung Hwang

## Abstract

Many of the biological functions performed by RNA are mediated by RNA-binding proteins (RBPs), and understanding the molecular basis of these interactions is fundamental to molecular biology. Here, we present MPRNA-immunoprecipitation (MPRNA-IP), an adaptation of the previously developed massively parallel RNA assay (MPRNA), and a new avenue for *in vivo* high-throughput dissection of RNA-protein interactions. By using custom pools of tens of thousands of RNA sequences containing systematically designed truncations and mutations, we are able to identify RNA domains, sequences, and secondary structures necessary and sufficient for protein binding in a single experiment. We show that this approach is successful for multiple RNAs of interest including NORAD, MS2, and human telomerase RNA, and we describe statistical models for identifying RNA domains and parsing the structural contributions of RNA in these interactions. By blending modern and classical approaches, MPRNA-IP provides a novel high-throughput way to elucidate RNA-based mechanisms behind RNA-protein interactions.

## INTRODUCTION

Noncoding roles of RNA have been known for decades in important biological processes. Many such RNAs exert biological roles by interacting with proteins. For example, both telomerase and the ribosome are ribonucleoprotein complexes with essential RNA components that both recruit essential protein subunits to the complex and perform integral roles in their respective biochemistry ^1,2^. This is true even within coding sequences, many of which contain regulatory elements that affect expression of the encoded protein. In such cases, RNAs bind to multiple proteins through specific and nonspecific elements, each of which can mediate different effects on the RNA’s function and/or metabolism ^3,4^.

While the biochemical versatility of RNA has made it a major player in many biological processes, this versatility complicates the dissection of underlying mechanisms. RNA not only uses primary nucleotide sequences but also secondary or tertiary structures for its biochemical activity (as in programmed −1 ribosomal frameshift signals ^5,6^), and often contains multiple functional domains (as in the lncRNA Xist ^7,8^). Therefore, one must consider multiple working modes or regions to understand even a single RNA’s mechanism of function, making the study of RNA technically difficult. Historically, these problems have been tackled through the use of truncation analysis, which breaks a molecule of interest into fragments to identify regions both necessary and sufficient for function. This can then be combined with more fine-grained approaches such as scanning mutagenesis or compensatory mutation analysis to home in on the specific sequences and structures required for a biological activity. However, comprehensively performing both truncation and mutation analyses with classical, low-throughput approaches is both time- and labor-intensive, ultimately slowing the pace of research on the mechanisms of noncoding or regulatory functions of RNA.

Previously, we and others developed a massively parallel RNA assay (MPRNA) that combines modern high-throughput oligonucleotide synthesis and next-generation sequencing to perform dissective analyses of RNA on a massive scale ^9,10^. This assay was used by us and others to study long noncoding RNA (lncRNA) sequences that drive nuclear localization. By tiling oligos across nuclear lncRNAs and testing whether each sequence could drive nuclear localization of an otherwise cytoplasmic construct, these studies were able to identify RNA elements that drive nuclear localization and sequence motifs enriched in these domains.

Here we advance MPRNA to interrogate *in vivo* RNA-protein interaction, combining it with immunoprecipitation (MPRNA-IP) and employing a complex tiling and mutagenesis approach to explore several key questions regarding RNA-based features of RNA-protein interactions: (1) whether an RNA can be divided into autonomous domains responsible for binding to an RBP, and (2) what RNA sequences and/or RNA secondary structures are necessary and sufficient for the RNA-protein interaction. We also make several technical improvements to increase the robustness of MPRNA, including the use of multiple molecular barcodes per RNA test sequence and the introduction of a new computational tool to analyze the sequencing outputs of MPRNA-IP. Using the examples of PUM2-binding RNAs, the MS2-MCP interaction, and human telomerase RNA-TERT interaction, we show that MPRNA-IP can identify protein-binding domains within an RNA, as well as the underlying RNA sequence motifs and RNA secondary structures driving RNA-protein interactions. Because RNA-protein interactions are essential for many noncoding roles of RNA, MPRNA-IP will serve as a powerful RNA-oriented high-throughput tool that is generally applicable to the study of regulatory RNA and will advance our understanding of RNA-protein interactions.

## RESULTS

### Overview of MPRNA-IP and the optimization of experimental parameters

MPRNA involves design of a custom pool of DNA oligonucleotides that are expressed and then tested in a given assay of interest *en masse*. While different custom pools were used for each of our MPRNA-IP experiments (Table 1), both the general anatomy of the oligonucleotides and the general experimental scheme remained constant. Each oligonucleotide includes 157 bases of experimental sequence (hereafter referred to as a tile) and a unique 10-mer molecular barcode, in addition to universal 5’ and 3’ PCR primer binding sites (Supplementary Figure 1A). Here, we use ‘oligo’ to mean the unique combination of tile sequence and barcode sequence. The steps of MPRNA-IP are summarized in Figure 1A – briefly, the oligo pools were cloned into an expression vector and transfected into HEK293T cells. Interaction of expressed oligos with the target protein of interest was assayed by formaldehyde-crosslinked RNA immunoprecipitation (fRIP) after an antibody was confirmed (Supplementary Figure 1B). The resulting RNA samples were processed for high-throughput sequencing through reverse transcription, PCR amplification of the recovered oligos, and generation of sequencing libraries of those amplicons. Finally, a generalized linear regression model was applied to every tile to identify oligos enriched in IP relative to total lysate (input), in which the numbers of sequencing reads were modeled by either Poisson or Negative binomial distributions and were compared between input and IP (see Methods).

**Figure 1:**
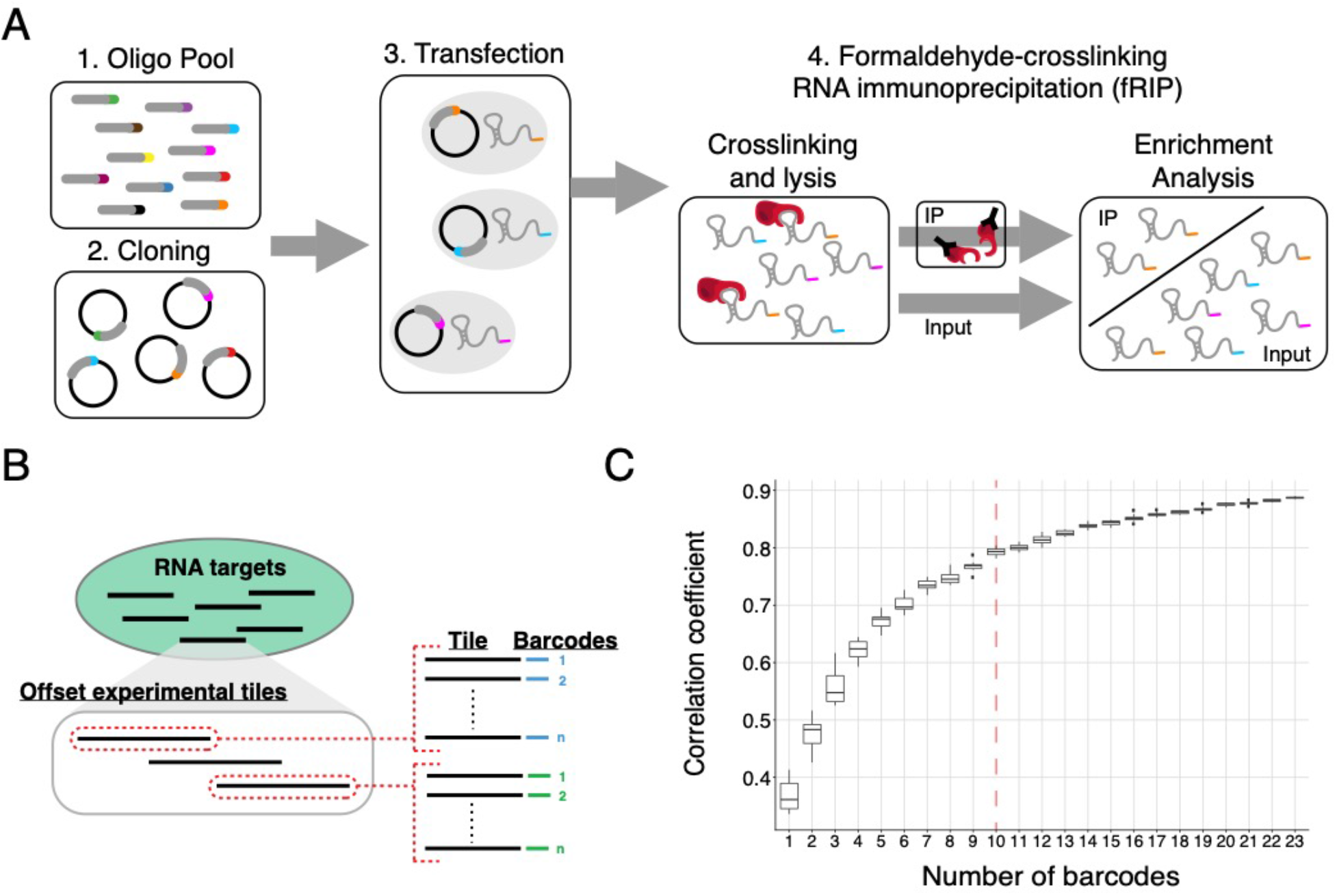
Massively parallel RNA assay to explore RNA-protein interactions. **(A)** MPRNA-IP assay. Oligonucleotide pools are cloned into an expression vector and transfected into HEK-293T cells. RNA tiles are assayed for protein interaction by formaldehyde-crosslinking RNA immunoprecipitation, to identify features that preferentially interact with a target protein. **(B)** To improve on previous versions of MPRNA, single experimental sequence tiles are represented by multiple unique barcodes. This strategy improves reliability and statistical power of the assay. **(C)** Optimization of MPRNA-IP experimental parameters. We simulated two biological replicates of MPR-NA-IP for a given barcode number by randomly selecting sequencing reads from a test experiment and determined the optimal number of barcodes required for high reproducibility of enrichment of tiles. 10 unique barcodes was sufficient to achieve >80% correlation between two biological replicates, with diminishing improvement above 10.

**Table 1.** Summary of oligo pools in the current manuscript.

| Oligo pool | Tile number of tiles | The number of barcodes per tile | Total number of oligos |
| --- | --- | --- | --- |
| **PUM2-binding RNAs** | 1,599 | 22 | 35,178 |
| **MS2-MCP** | 105 | 50 | 5,250 |
| **hTR-TERT** | 614 | 25 | 15,350 |

To improve on previous implementations of MPRNA, we assigned multiple unique molecular barcodes to each experimental tile, adding internal technical replicates to the experimental design (Figure 1B). The purpose of this redundant design was to (1) optimize the representation of oligos, (2) reduce the likelihood of false positive results driven by an individual barcode, and (3) increase statistical power in enrichment analysis. To determine the optimal number of barcodes for each experimental sequence, we randomly down-sampled sequencing reads from an MPRNA-IP experiment (hTR-TERT in Table 1) and generated simulated replicate pairs for a wide ranges of barcode numbers per tile. Then we calculated Pearson correlation coefficients of average enrichment (IP/input) across tiles between the replicates. Higher numbers of molecular barcodes drove higher correlation between the two simulated replicates (Figure 1C), and 10 molecular barcodes was sufficient to achieve a correlation of 0.8. Thus, moving forward with new oligo pool designs, we included at least 10 unique barcodes for every tile.

Next, we optimized sequencing depth to guide cost-effective sequencing of MPRNA-IP samples. Here, we again randomly down-sampled sequencing reads of an MPRNA-IP experiment (hTR-TERT in Table 1) and observed the number of missing oligos (oligos that were not detected in input samples) at various simulated sequencing depths. As the number of sequencing reads in high-throughput sequencing is known to depend on GC percentages, we evaluated 100 random sequences that have roughly 50% GC content, allowing unbiased calculation of the drop-out rates. Each random sequence had 25 molecular barcodes, making for 2,500 oligos in total. As expected, the proportions of missing oligos decreases as the average sequencing depth per oligo increases (Supplementary Figure 1C). We determined that at least 700 sequencing reads per oligo is recommended to minimize the number of missing oligos.

### MPRNA-IP can recover protein-binding RNA domains and underlying sequence motifs

One of the mechanisms driving specificity for RNA-protein interactions is protein recognition of an RNA primary sequence motif. To test whether MPRNA-IP can successfully probe primary sequence elements, we explored Pumilio (PUM)-binding RNAs, as the Pumilio-RNA interaction has been studied in great detail ^11^. To design the experimental tiles within this MPRNA oligo pool, we selected 23 PUM2-binding RNAs from those identified in a previous study via enhanced crosslinking and immunoprecipitation (eCLIP), a technique for identifying the location of protein binding within endogenous RNAs^12^, covering a broad range of number and density of eCLIP peaks per molecule (Supplementary Figure 2A). We tiled the entire length of these RNAs with overlapping fragments of 157 nucleotides to give a total of 1599 experimental sequences. Each tile is represented by 22 molecular barcodes, generating a final pool of 35,178 oligos. MPRNA-IP was performed as in Figure 1A, using an antibody raised against PUM2 for immunoprecipitation.

We first confirmed that the oligo pool was well-represented in our assay conditions by assessing what percentage of oligos in the pool were detected above a given threshold in our input sample. Across three biological replicates, about 90% of oligos were detected in the input sample (Supplementary Figure 2B) and the distributions of reads per million (RPM) values of the oligos were comparable (Supplementary Figure 2C). Importantly, enrichment of the tiles was highly correlated across biological replicates (Supplementary Figure 3).

Next, we identified the statistically enriched tiles in the anti-PUM2 IP samples using a generalized linear regression model comparing the relative number of sequencing reads between the input and the IP per a tile while controlling for the effect of biological replicates and molecular barcodes (see Methods). 155 of the 1599 experimental tiles were significantly enriched in the IP relative to the input (False Discovery Rate (FDR) < 0.05 and IP/input>1, Supplementary Figure 4), and we saw a high overlap between these enriched tiles and eCLIP peaks, as shown for the representative PUM2-binding lncRNA NORAD (Figure 2A). Also, the number of enriched tiles correlated with the number of eCLIP peaks across multiple RNA targets (Figure 2B, n=23, Pearson correlation=0.7, p-value=0.0002191), suggesting that RNA-PUM2 interaction is mediated by local RNA elements that can be identified by the tiling approach of MPRNA-IP. These data demonstrate that MPRNA-IP can identify distinct, autonomous, and biologically relevant regions of these RNAs sufficient for binding to PUM2, independent of the full-length RNA.

**Figure 2:**
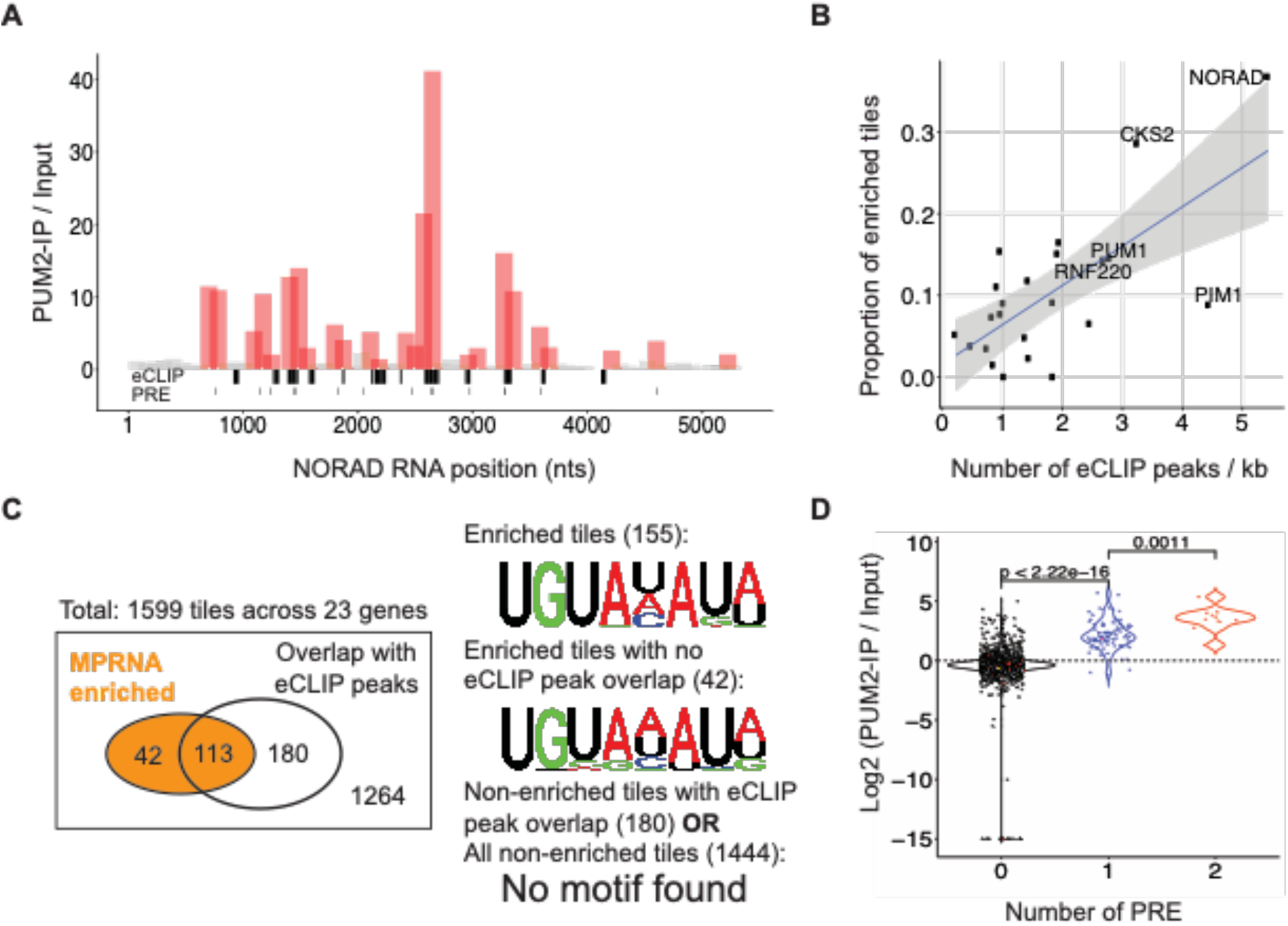
MPRNA-IP can identify regions and primary sequence motifs driving specificity of pumilio proteins. **(A)** MPRNA-IP tiles are described on the well-known pumilio binding RNA NORAD. Red tiles are statistically enriched in the IP sample and show high correspondence with both PUM2 eCLIP peaks and the known PUM2-binding consensus motif, the pumilio responsive element (PRE) **(B)** Correlation between the number of eCLIP peaks within a target RNA and the proportion of tiles enriched by MPRNA-IP. **(C)** Motif analyses of tiles enriched by MPRNA-IP confirms the well-known pumilio responsive element (#155). Tiles enriched by MPRNAthat do not overlap with previously generated eCLIP peaks again produce PRE (#42). Tiles within an eCLIP peak that were not also enriched in MPRNA-IP (#180) did not retrieve any consensus motif. (D) Enrichment of a tile correlates with the number of PREs contained within that tile, indicating a causal role of the PRE in tile enrichment.

We, furthermore, explored if MPRNA-IP can accurately determine RNA motifs bound by a given protein. Previous studies have identified the Pumilio Response Element (PRE) as the RNA sequence motif that drives interaction with Pumilio proteins ^13,14^. Unbiased motif analysis of the enriched tiles using the motif discovery tool (MEME) successfully recovered the PRE (Figure 2C). Moreover, we found that the tiles that were not enriched do not have any specific sequence motif, suggesting a low false negative rate. Finally, we compared our MPRNA enriched tiles with the known peaks identified by eCLIP that were often used to identify RNA elements. Out of the 155 enriched tiles, 42 did not overlap with eCLIP peaks, but motif analysis showed that these tiles still contained the PRE. Contrastingly, when we performed motif analysis on tiles that were not enriched by MPRNA-IP, even those that overlap with eCLIP peaks, the PRE was not found. This implies that eCLIP may not have the sensitivity to recover binding at these regions, or perhaps that these regions are hidden from PUM2 in the full-length RNA that eCLIP deals with. MPRNA-IP results further showed that the enrichment strength of the tiles is correlated with the number of PREs, revealing the causal effect of the PRE in this interaction (Figure 2D). These results show that MPRNA-IP can identify a primary sequence motif underlying an RNA-protein interaction at least comparably to a full-length RNA-based approach like eCLIP.

### Mapping functional RNA structures through protein binding

In addition to proteins that recognize primary RNA sequence, decades of research have shown that RNA secondary structures drive many RNA-protein interactions ^15–17^. To determine MPRNA-IP’s ability to identify the contribution of structure to RNA-protein interaction, we expanded our oligo pool design strategy to incorporate classical compensatory mutation analysis into MPRNA-IP (Figure 3A). Here, we first predicted all potential stems of at least 2 uninterrupted base-pairs based on the base-pairing rules of RNA primary sequence (A:U, G:C, G:U). Next, the 5’ and 3’ sides of a predicted base-paired structure were individually mutated to their complementary sequences to disrupt the predicted stem, and then mutated together in a single compensatory mutant to restore putative base pairing (e.g., a CU-AG stem would be mutated to GA-AG and CU-UC to disrupt and to AG-UC to restore). In these experiments, when both the 5’ and 3’ mutants impair biochemical function and are partially or fully rescued by the compensatory mutant, this serves as strong evidence that the predicted structure forms and is important for the function being tested (in this case, protein binding). After performing MPRNA-IP experiment, we applied a regression-based computational model that considers differential enrichment of wild-type (WT), mutant, and compensatory mutant sequences to quantify the effect of base pairing within a stem on an RNA-protein interaction, independent of the effects of changing the primary RNA sequence. Specifically, this regression model assumes that the difference in enrichment between WT and the 5’ or 3’ mutant is a sum of both sequence and base-pairing effects, while the enrichment difference between the compensatory mutant and the 5’ or 3’ mutant is due only to base-pairing effects (the bottom panel in Figure 3A and see Methods). We refer to the estimated base-pairing effect as “stem score”.

**Figure 3:**
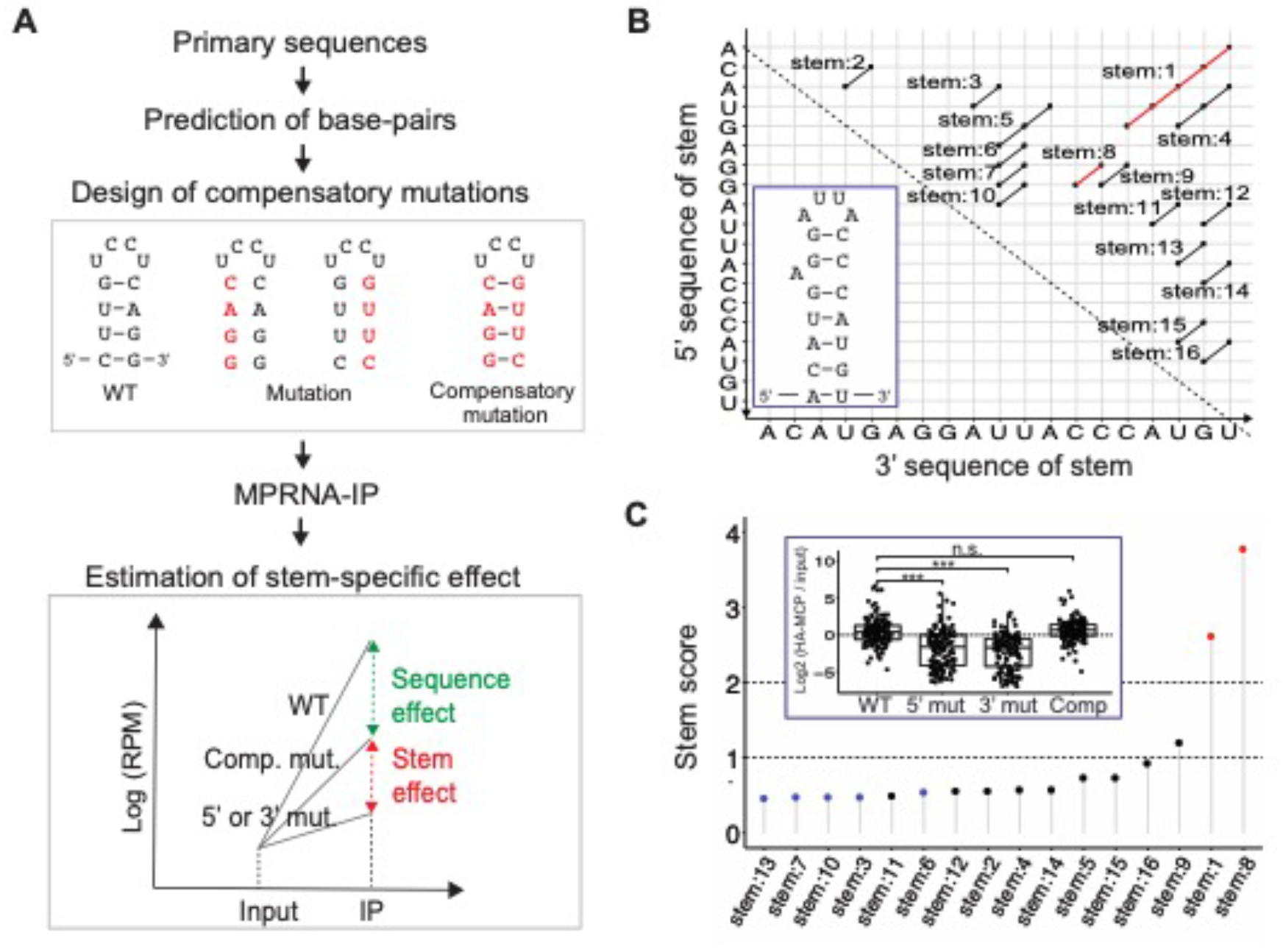
Structural features of the MS2 hairpin are identified by MPRNA-IP. **(A)** Schematic demonstrating compensatory mutation analysis to explore RNA structure by MPRNA-IP. After predicting secondary structure from primary sequence, point mutations or full-stem mutations were introduced into the 5’ or 3’ ends of the predicted structure, as well as both 5’ and 3’ mutations in conjunction to rescue the putative structural element. A regression model is applied to parse the structural and sequence-based contributions to binding, based on compensatory mutation analysis. **(B)** All possible stems containing two or more adjacent base pairs, based on the primary sequence of the MS2 hairpin. The two known stems within the wildtype MS2 hairpin are marked in red. The inset represents the known structure of the MS2 hairpin. **(C)** Stem score of all predicted base-pairs in (B), estimated by the statistical model presented in (A). The top stem score is observed for stems 1 and 8, the two stems known to form in the wildtype hairpin. Inset: distribution of oligos for stem 8. 5’ and 3’ mutation of stem 8 disrupts interaction with MCP, which is rescued by compensatory mutation to regenerate the stem. Data points represent 50 barcodes across three biological replicates of MPRNA-IP.

We chose as a model the well-studied bacteriophage MS2 RNA hairpin (Figure 3B inset) that is critical for binding to the MS2 coat protein (MCP). After predicting all 16 potential stems of at least 2 uninterrupted base-pairs based on the MS2 primary sequence (Figure 3B), we synthesized a pool of 5,250 oligos covering 5’, 3’ and compensatory mutations for all the predicted base-pairs. Then we co-transfected our oligo pool with an HA-FLAG-tagged tandem dimer of MCP and performed immunoprecipitation using an anti-HA antibody (Supplementary Figures 1B, 5A and 5B). Our MPRNA-IP results showed that, among all the possible stems, “stem:1” and “stem:8”, corresponding to the two known stems of the MS2 hairpin, have the highest stem scores on MCP binding (Figure 3C). Considering as an example stem:8 (corresponding to the GG-CC stem directly below the apical loop of the hairpin), tiles containing either 5’ or 3’ mutations were significantly depleted relative to wild type, while compensatory mutation restored the enrichment comparable to wild type levels (Figure 3C inset). These results demonstrate that MPRNA-IP can successfully map functional RNA secondary structures through protein-binding, expanding the utility of MPRNA-IP beyond probing primary sequence only.

In addition to mutating predicted stems, we included tiles containing point mutations at every nucleotide position to better assess the contribution of primary sequence to binding. Our results showed that most point mutations decrease enrichment to varying degrees, but a U11C or U11A mutation (the −5 position relative to the MS2 RNA translational start) strengthened the binding (Supplementary Figure 5C). Although the increased enrichment by these mutations lacks statistical significance (FDR<0.05) potentially due to the small magnitude of perturbations, it is consistent with previous findings that mutation of this nucleotide to C enhances MS2-MCP interaction ^18,19^.

### Application of MPRNA-IP to human telomerase RNA

After validating the ability of MPRNA-IP to identify RNA domains, sequence motifs, and structures that mediate protein interaction, we sought a new experimental system to apply all functionalities of this assay. Telomerase RNA (TR) is an essential ncRNA component of telomerase, both serving as a scaffold for the RNP and providing the template for reverse transcription that is required for telomere lengthening ^20,21^. Although TRs are diverse in primary sequence and length across species from yeast to human, all vertebrate TRs contain three conserved RNA domains: template/pseudoknot, CR4/5, and H/ACA domains (Figure 4A) ^22^. Both CR4/5 and the template/pseudoknot domains have been shown to interact with the telomerase reverse transcriptase (TERT) protein, the catalytic subunit of telomerase, but as two separate modules ^23,24^. However, it remains unclear precisely which sequence and structural elements within human TR (hTR) are required for TERT-binding. Therefore, we explored the hTR-TERT interaction as a biological example to test how multiple aspects of MPRNA-IP can be used to study the RNA features involved in a biologically important RNA-protein interaction. As hTR is 451 nucleotides and our oligos are ~200 long, we divided it into three fragments (tiles), covering the three known domains (Figure 4A). In addition to WT sequences, we included tiles containing either deletions or disrupting and compensatory mutations of all stems within the known hTR secondary structure. These experimental tiles are accompanied by 100 tiles of random sequences to serve as negative controls in statistical calculations. Each experimental and control tile is represented by 25 barcodes (Supplementary Figure 6A).

**Figure 4:**
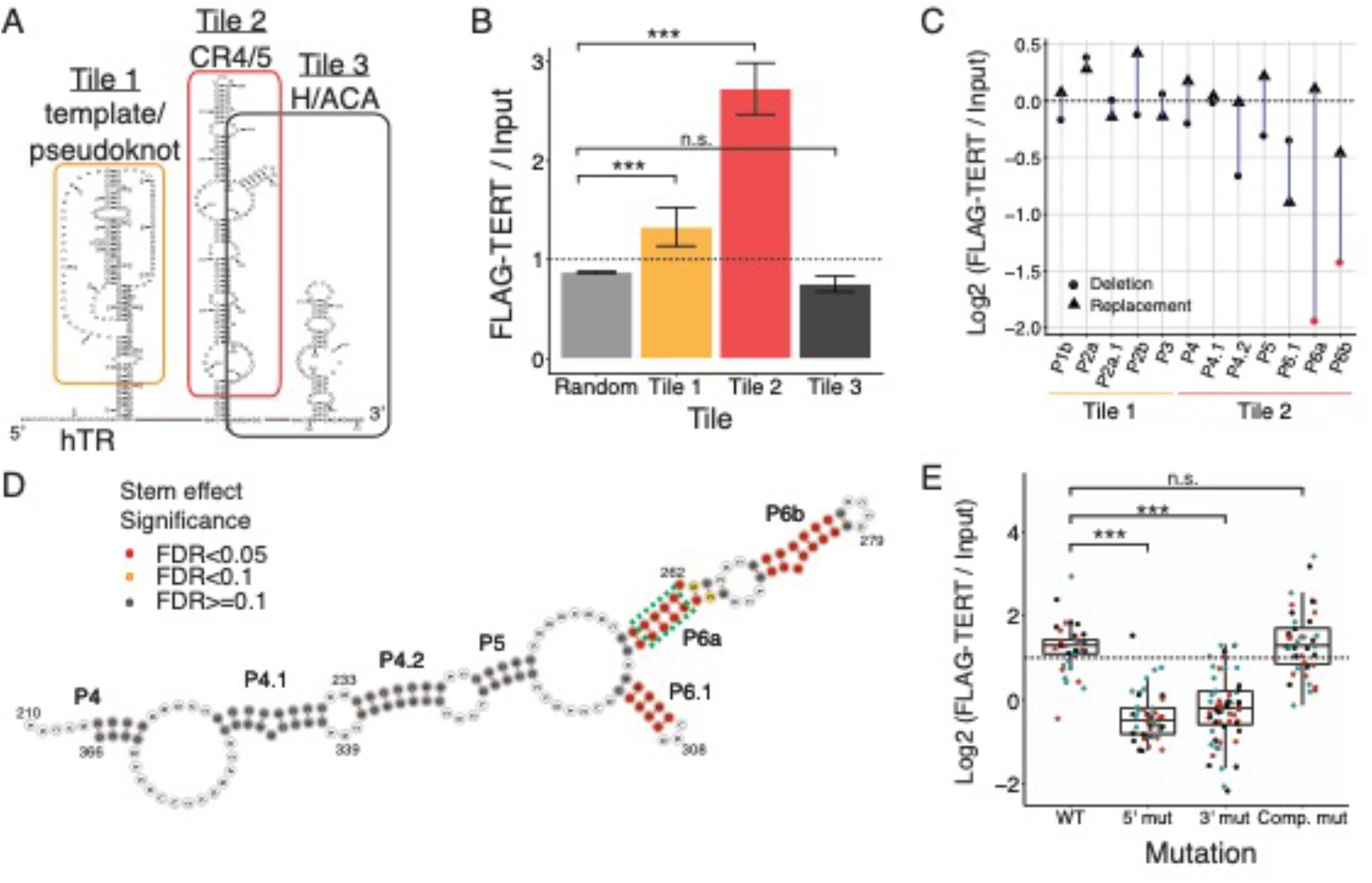
Comprehensive application of MPRNA-IP to explore domains, sequence, and structural contributions of RNA in human telomerase RNA-TERT interaction. **(A)** Schematic of the known structure of human telomerase RNA (hTR). For MPRNA pool design, hTR was divided into three tiles covering the pseudoknot (tile 1, orange), CR4/CR5 domain (tile 2, red), and snoRNA-like domain (tile 3, black). The approximate RNA locations where TERT binds are shown. **(B)** Enrichment of oligos, measured by fold change of RPMs in FLAG·TERT relative to Input, were summarized according to the comparing tiles. The 3 biological replicates with each multiple oligos were pooled to compare mean enrichments across tiles **(C)** The changes of enrichment in deletion (denoted by circle) and replacement (denoted by triangle) relative to wild type sequences are described. Significant changes (FDR<0.05) were indicated by red. **(D)** Statistical significances of stem scores of all known stems in tile 2 were overlaid on tile 2’s secondary structure. The most signifnicant p-value is used for a given location if there are multiple tiles covering the location. The stem with the largest stem score is denoted by a dotted box in P6a. **(E)** The enrichments of oligos of the tile corresponding to the dotted box in (D) are described according to mutations. 3 biological replicates are denoted by different colors.

Using this pool of hTR sequences and a FLAG-TERT IP, we first tested if MPRNA-IP could selectively enrich for the hTR domains known to bind TERT. The two tiles covering the template/pseudoknot and CR4/5 domains were significantly enriched in the IP relative input (FDR<0.05), while both the H/ACA domain tile and the 100 random sequences were not (Figure 4B and Supplementary Figure 6B), consistent with previous reports showing that pseudoknot and CR4/5 domains bind to TERT independently ^23,25^. Moreover, the tile covering the CR4/5 domain is enriched more highly than that covering the template/pseudoknot domain (2.38 vs. 1.26 by the fold change of IP to Input). This may imply that the CR4/5 domain is the primary TERT-interaction module, as has been previously suggested^23,25^, though more direct experimental evidence will be needed to confirm this finding.

Next, we attempted to identify secondary structures within the template/pseudoknot and CR4/5 regions required for binding to TERT. We first compared oligo enrichment between wildtype tiles and tiles with individual stem deletions (see Methods). Overall, deletion of stems in CR4/5 caused a loss of enrichment (mean enrichment: 0.68) while deletion of stems in the pseudoknot did not show significant effects (mean enrichment: 1.04) (Figure 4C). Furthermore, replacing the deleted P6a and P6b stems with their compensatory mutant sequences to reform the original structures recovered TERT enrichment, indicating that the secondary structures are necessary for TERT binding.

To more rigorously dissect the RNA structural contributions to hTR-TERT interaction, we expanded our mutation strategy to include compensatory mutation analysis in both 1-base-pair and 4-base-pair sliding windows across all known stems and calculated stem scores to quantify the effect that base-pairing has on binding. We identified 10 tiles showing significant stem scores (Supplementary Table 1 and Supplementary Figures 7A and 8A), all of which are located in stems P6a, P6b, and P6.1 within the CR4/5 domain (Figure 4D and Supplementary Figure 7). Specifically, 5’ and 3’ mutants of stems P6a, P6b, and P6.1 showed reduced enrichment that was rescued by compensatory mutation (Figure 4E and Supplementary Figures 7B) while the other stems did not show this pattern (for example stem P5, see Supplementary Figures 7C). In contrast to the CR4/5 domain, we did not find any significant stem effects in the template/pseudoknot domain, despite the known interaction between TERT and the pseudoknot (Supplementary Figure 7A, stems P1b, P2a.1, P2a, P2b, and P3). This suggests that pairing alone in these regions is not sufficient for binding by TERT and that specific sequences within these stems may also be required for the interaction. Together, these results demonstrate that MPRNA-IP can be used to rigorously explore the RNA elements required for protein binding, all in a single experiment.

## Discussion

RNA-protein interactions play central roles in many biological processes, and modern high-throughput assays have been used by many labs to expand our understanding of these interactions. Approaches such as RNA IP (RIP) and crosslinking and immunoprecipitation (CLIP) have proved invaluable, and advancements in various CLIP-based sequencing approaches, including HITS-CLIP ^26^, PAR-CLIP ^14^, iCLIP ^27^, and eCLIP ^12^, have successfully identified RNAs interacting with RBPs, providing insights into associated diseases and development. However, CLIP-based methods are limited in how much they can reveal about the molecular principles governing RNA-protein interactions from the perspective of RNA, as these techniques cannot directly test whether an RNA element or sequence motif is necessary or sufficient for the interaction. Here we describe MPRNA-IP, an *in vivo*, high-throughput sequencing-based approach for dissecting the features of RNA, including domains, sequence motifs, and stem structures, that are necessary and sufficient for protein binding.

In our first test of this approach, we employed a tiling strategy to scan across 23 RNAs known to bind to the well-characterized RNA-binding protein PUM2 ^28^. We observed a strong enrichment of tiles containing the known consensus motif for PUM2 binding, and enriched tiles showed a high degree of overlap with peaks from previously published PUM2 CLIP data ^12^. However, our experiment displayed a few important differences from CLIP. Firstly, we observed enrichment of several tiles that contained the PUM2-binding consensus but did not overlap with a PUM2 CLIP peak. This is intriguing, because it represents RNA domains that are capable of binding PUM2, harbor the PRE motif, yet are not detectably bound when assayed by CLIP. Perhaps, these domains are masked from PUM2 binding by secondary structure or by binding of another RBP. Nonetheless, comparison with eCLIP data demonstrates that MPRNA-IP can accurately and faithfully capture the *in vivo* requirements for RNA-protein binding. Secondly, by isolating a small region of an RNA from its native context of the full-length molecule, we are able to test what elements are sufficient for protein-binding independent of the rest of the RNA. This concept of sufficiency cannot be tested with approaches that survey the full-length endogenous transcriptome but is key in understanding any biological activity including protein binding.

In our second test of the MPRNA-IP approach, we used a more complicated dissection strategy – that of compensatory mutation analysis – to isolate the secondary structure elements of the bacteriophage MS2 hairpin that are required for binding to its partner protein MCP. Importantly, in an attempt to accurately simulate a situation in which nothing is known about the target RNA’s structure, we performed compensatory mutational analysis on all possible stems of 2 base pairs or longer in the MS2 hairpin sequence (i.e., not just the known structures). By computationally separating the effect that base-pairing has on protein binding from the effect of sequence changes, we were able to clearly distinguish the two true stems of the MS2 hairpin from others that were predicted. This is a significant advancement in the application of compensatory mutation analysis, which has traditionally only been used to test potential stems one-by-one. Furthermore, this approach to dissecting secondary structures in the direct context of biological function is likely to be especially advantageous in the study of long noncoding RNAs. lncRNAs show low conservation of primary sequence, and it has been speculated that structural conservation actually drives conserved function ^29–31^. Traditional chemical probing methods ^32^ have been successfully used to determine lncRNA structure, but extensive validation is often required to determine whether or not the resulting structural models are functionally relevant. This MPRNA-IP approach avoids this problem by directly analyzing the effects of potential secondary structure disruption and restoration on protein binding, effectively identifying structures through their biological function.

Having successfully employed tiling and compensatory mutation strategies in MPRNA-IP to identify known protein-binding RNA elements, we next applied these approaches to the interaction between the protein TERT and the human telomerase RNA (hTR), an RNA-protein interaction with mechanisms that are not fully understood. While the structural domains of hTR required for TERT binding were identified 15-20 years ago (the template/pseudoknot and CR4/5 regions) ^23,33^, it has remained unclear which specific structural elements within those domains are required for binding to TERT. Using a tiling approach to split the RNA into separate domains and applying compensatory mutation analysis to the known stems of the RNA, we were able to identify regions within the CR4/5 domain where base-pairing is required for binding to TERT. Our *in vivo* results identifying P6a, P6b, and P6.1 as TERT-binding stems are consistent with previous *in vitro* results identifying a minimal CR4/5 domain that is sufficient for telomerase activity ^34^.

Unlike the CR4/5 domain, we did not find any stems within the template-pseudoknot domain of hTR in which restoring base-pairing via a compensatory mutant restored TERT binding, but this is not necessarily unexpected. The pseudoknot of hTR is a rather intricate secondary structure and is located very close to the catalytic center of the telomerase enzyme ^35^. Therefore, it seems likely that both base-pairing and specific sequences within the pseudoknot are required for binding to TERT. A previous study also did not find substantial inhibition of TERT interaction by the mutations in the pseudoknot domain, so it may also be the case that larger mutations or deletions are required to fully disrupt TERT-binding to the pseudoknot ^25^. Furthermore, it is perhaps not surprising that strong stem effects would be relatively rare, not just in hTR but in all RNAs. Many large RNAs are known to contain long linker regions in between their functional protein-binding sites (e.g., NORAD ^36,37^, yeast telomerase RNA ^21,38^, Xist ^8^), and many RNA-binding proteins have preferences specific to sequence, not secondary structure ^39^. Therefore, it follows that regions where base-pairing alone is required for protein binding would be relatively uncommon. This complexity in the requirements for protein binding further emphasizes the advantage of using customized sequence pools in these MPRNA-IP experiments. While compensatory mutation analysis is the most complex mutational strategy employed in this study, even more complex strategies could be employed in the future to dissect interactions in which both sequence and structure are required or including internal deletions to assess long-range interactions.

Many methods have been developed previously that, like MPRNA-IP, use a large pool of RNA sequences to test the requirements for protein binding, including RNAcompete ^40^, RNA-RAP ^41^, RNA-MaP ^42^, RBNS ^43^ and HiTS-KIN ^44^. However, MPRNA-IP differs from these approaches in two key aspects. First, while all of these previous approaches are performed *in vitro*, MPRNA-IP is performed *in vivo*, allowing for these biochemical analyses to be performed under the most physiologically relevant conditions possible. Additionally, this allows for biochemical analysis of RNA-binding preferences even for proteins that are resistant to biochemical purification, proteins for which *in vitro* approaches would be inviable. Secondly, the test sequences used in MPRNA-IP are markedly larger than those used in most previous high-throughput dissection approaches. As demonstrated by our experiments with human telomerase RNA, this advancement allows for dissection of RNA domains such as CR4/5 that would be too large for other similar approaches, as well as in-depth characterization of base-paired regions. Despite the strengths of MPRNA-IP compared to similar assays, there are important considerations in implementing MPRNA-IP to a new system. First, many RBPs form complexes to bind to RNA and therefore stoichiometry should be considered when an overexpressed protein is used in MPRNA-IP. Second, previous studies observed cell-type-specific activities in high-throughput reporter assays, highlighting the importance of choosing an appropriate *in vivo* model system^45,46^. Finally, the degree of enrichment difference between input and IP varies depending on RNA-protein interactions, requiring proper control tiles in an oligo pool. With these parameters tuned for a context, MPRNA-IP will serve as a powerful systematic tool probing *in vivo* RNA-protein interactions in a high-throughput and quantitative way.

## METHODS

### Oligo Pool Design and Cloning

Each oligonucleotide includes 157 bases of experimental sequence (experimental sequences were chosen according to the unique goals of the MPRNA assay) and a unique 10-mer molecular barcode, in addition to universal 5’ and 3’ PCR primer binding sites (Supplementary Figure 1A). Designed oligo pools were synthesized by Twist Biosciences (San Francisco, CA). Cloning of oligo pools was performed according to previous MPRNA protocols (Shukla et al 2018). Briefly, oligo pools were amplified by emulsion PCR (Chimerx) to avoid sequence-based biases in amplification. ePCR products were purified, digested, then ligated into an expression vector. The transformation was done with Top10 chemically competent cells (Invitrogen). After harvesting the cells, plasmid DNA was isolated. The representation of the pool was verified after ePCR amplification and after plasmid isolation by Illumina sequencing.

### MPRNA-IP

An oligo pool was transfected to HEK293T using the X-tremeGENE HP DNA transfection reagent (Millipore Sigma) according to manufacturer’s instructions. Interaction with the target protein of interest was assayed by formaldehyde-crosslinked RNA immunoprecipitation (fRIP) as previously described ^47^. Following immunoprecipitation and reverse crosslinking, RNA samples were reverse transcribed and amplified by PCR using primers containing unique sequencing identifiers that allowed for multiplexed sequencing. Amplicon sequencing was performed on an Illumina NextSeq. The sequencing reads of input and IP samples were assigned to the designed tiles by mapping barcode sequences. We also checked the sequence identity of up to 152 nts (limited by read size of our single end sequencing) between a sequencing read and a mapped tile and filtered the sequencing reads if they have more than 2 mismatches.

### Identification of enriched tiles in IP relative to Input

The numbers of sequencing reads assigned to a tile were compared between input and IP for an enrichment of a tile by applying a Poisson regression model. Specifically, the following model was constructed for a tile:

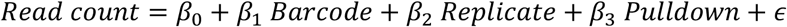

where an error term, *ϵ* follows a Poisson distribution; the categorical variables *Barcode* and *Replicate* describe unique barcodes and biological replicates, respectively; and the categorical variable *Pulldown* has two levels of Input and IP. In the model, the coefficient *β*_3_ represents the enrichment of a tile as a fold change (log scale) of IP relative to input while controlling barcode-specific and biological replicate-specific effects on enrichment. We applied this regression model to all the tiles that had at least 3 expressed barcodes in input samples. The barcodes were defined as expressed if their reads per million (RPM) values were greater than or equal to 1. The statistical significance of enrichment was evaluated by false discovery rate (FDR) that was computed based on an empirical estimation of null distribution. The R packages MASS and fdrtools were used for regression and empirical FDR calculation, respectively.

### Statistical model of compensatory mutation

For a potential or known stem, the following regression model is constructed to account for the variability of sequencing read counts across oligoes testing WT, mutation (either 5’ or 3’), and compensatory mutation:

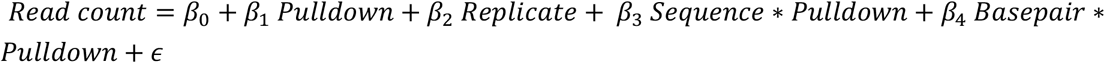

where error follows a negative binomial distribution, the categorical variables *Replicate* describe unique biological replicates, and the categorical variable *Pulldown* has two levels of total lysates and IP. The binary variables *Sequence* and *Basepair* represent two driving components of RNA-protein interaction and indicate whether they contribute to an oligo’s enrichment in IP relative to total lysate. We assume that WT has both sequence and base-pairing effects while compensatory mutation has base-pairing effects only, relative to 5’ or 3’ mutation. The R package MASS was used to fit the generalized regression model. Multiple test correction was done by using Benjamini-Hochberg’s FDR calculation.

### PUM2 MPRNA-IP analyses

A list of eCLIP peak regions of PUM2 was obtained from ENCODE (Accession ID: ENCFF880MWQ). Association of PRE with a tile is based on exact match of UGUAHAUA in the tile sequence, where H indicates any base other than G. Motif analyses was done by MEME.

## CONTRIBUTION

T.H. and J.L.R. conceived and directed the study. T.H., E.P.H., Y.H.L, and W.C. designed and performed experiments. Y.K.L., E.L., and J. S. S. assisted experiments. T.H. developed computational methods and performed statistical analyses. T.H., E.P.H., W.C., Y.H.L., E.L., and J.L.R. wrote the manuscript.

## ACKNOWLEDGEMENT

We are grateful to Joo Heon Shin for providing sequencing resources. We acknowledge the BioFrontiers Computing Core at the University of Colorado Boulder for providing High Performance Computing resources supported by BioFrontiers IT. This research was supported by NIH/NIGMS grant R00 GM137072 to T.H.

