## Supplemental File for "Massively Parallel Dissection of RNA in RNA-protein interactions *in vivo*"

**Supplementary Figure 1. Design of oligonucleotide pool and the optimization of MPRNA-IP.**

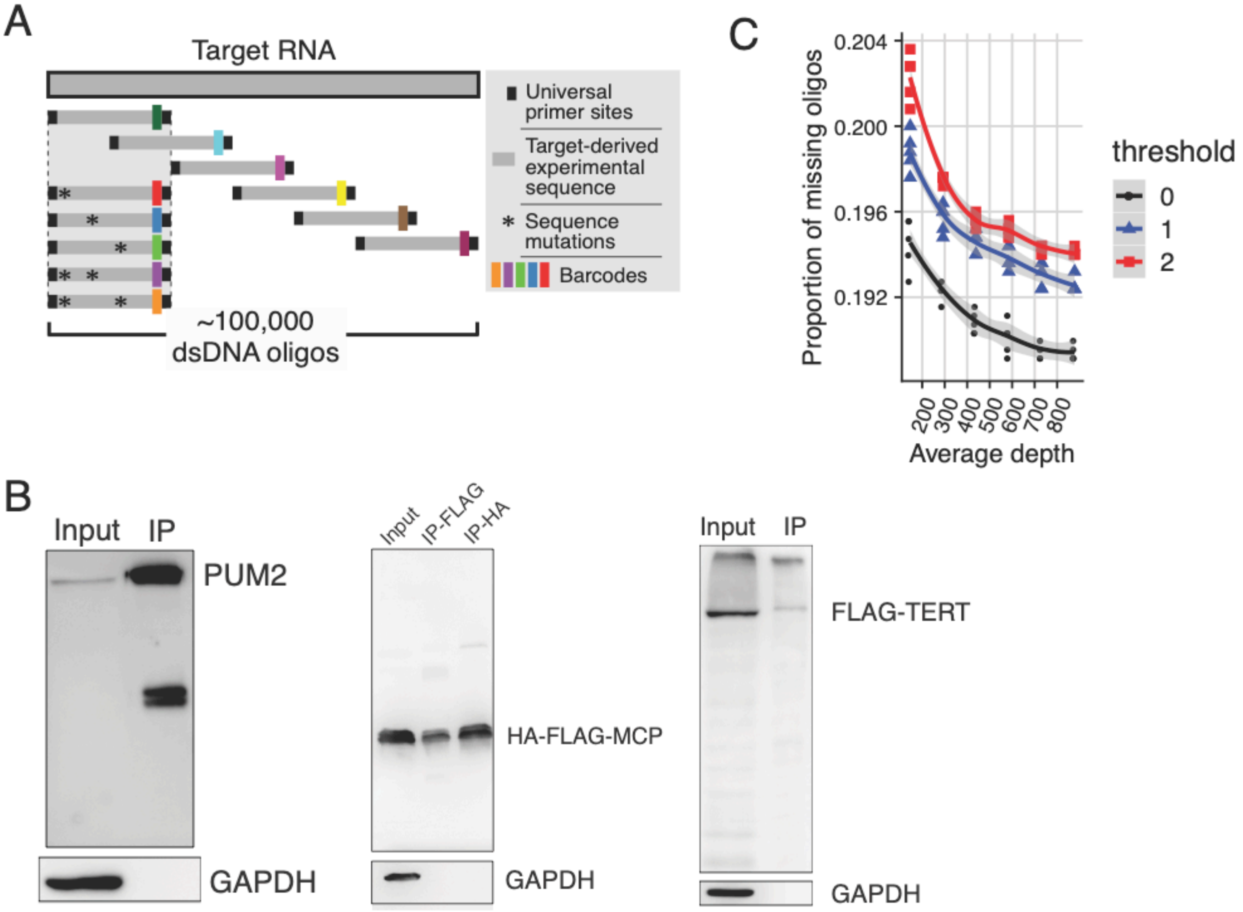


**(A)** Oligonucleotide pool design. Each unique oligo contains 157 bases of experimental sequence, a unique 10-mer molecular barcode, and universal 5' and 3' primers. The experimental sequences are derived from target RNAs and are offset but overlapping to minimize the missing of local RNA structures from truncation. Tiles also incorporate a strategic mutagenesis strategy to test sequence and the structure driving RNA-protein interaction. **(B)** Western blots showing efficient immunoprecipitation of target proteins for each MPRNA-IP experiment. For MS2-MCP MPRNA-IP, the MCP construct was both FLAG- and HA-tagged. As the IP against the HA tag showed higher correlations of enrichment across biological replicates compared to FLAG-IP, we focused on HA-tagged MCP for further analyses (Supplementary Figure 5B). **(C)** Optimization of sequencing depth for MPRNA-IP. About 700 sequencing reads/oligo are recommended to attain lower proportion of mission oligos to the total number of oligos in a pool. Here, "missing" oligo is defined as an oligo whose sequencing reads are lower than or equal to a given threshold value. Three different threshold values were used as indicated by different color.

**Supplementary Figure 2 MPRNA-IP for PUM2-binding RNAs.**

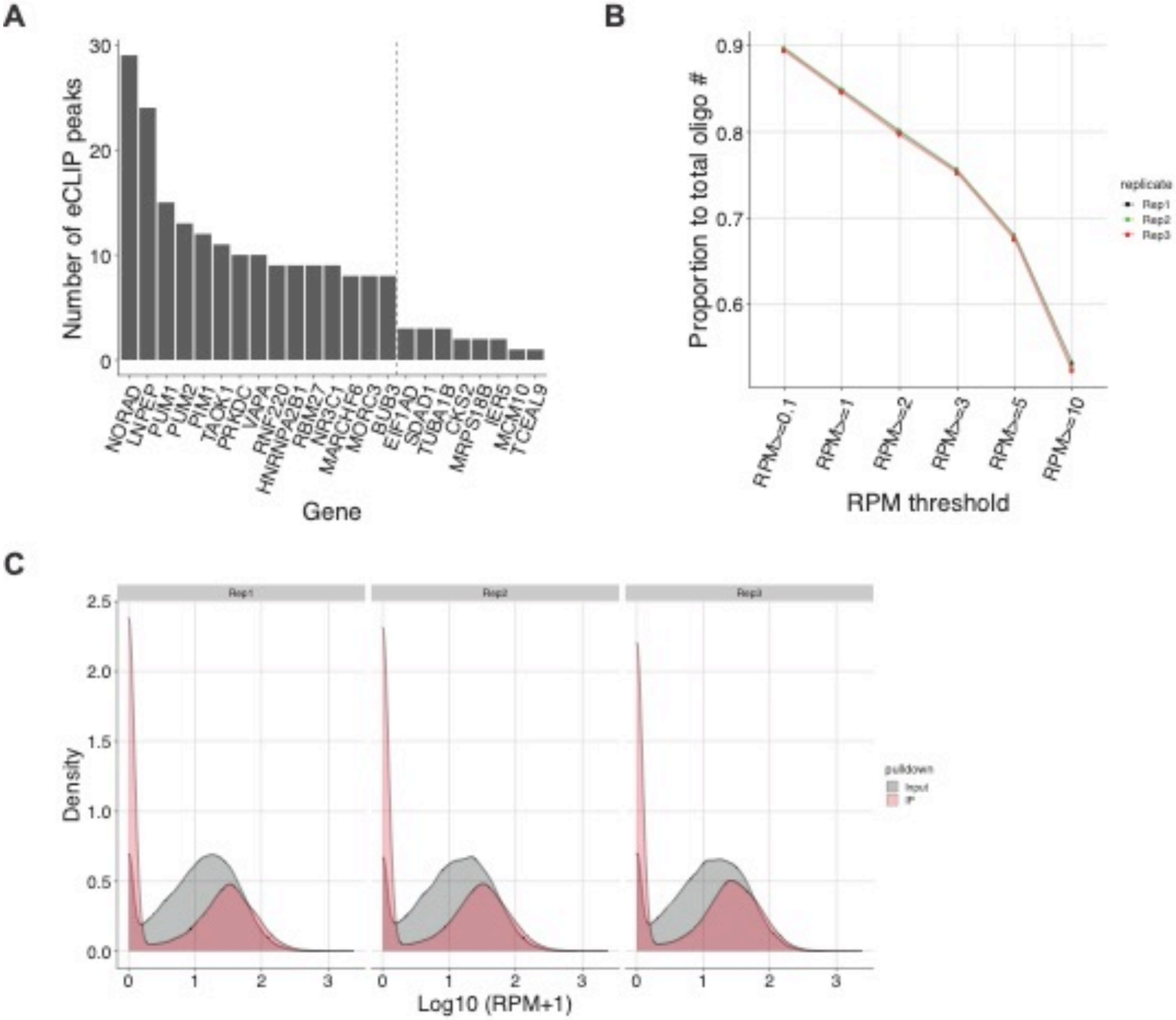


**(A)** RNA targets used for experimental tile design in the PUM2 MPRNA-IP pool. These RNAs were selected based on a previous eCLIP result, such that the pool contains multiple PUM2 targets covering diverse range of eCLIP peaks. **(B)** About 90% of designed oligos have an RPM $\geq$ 0.1 and are highly reproducible between replicates. **(C)** The distribution of RPM across oligos in input and IP are described for three biological replicates.

**Supplementary Figure 3. Reproducibility of MPRNA-IP for PUM2-binding RNAs.**

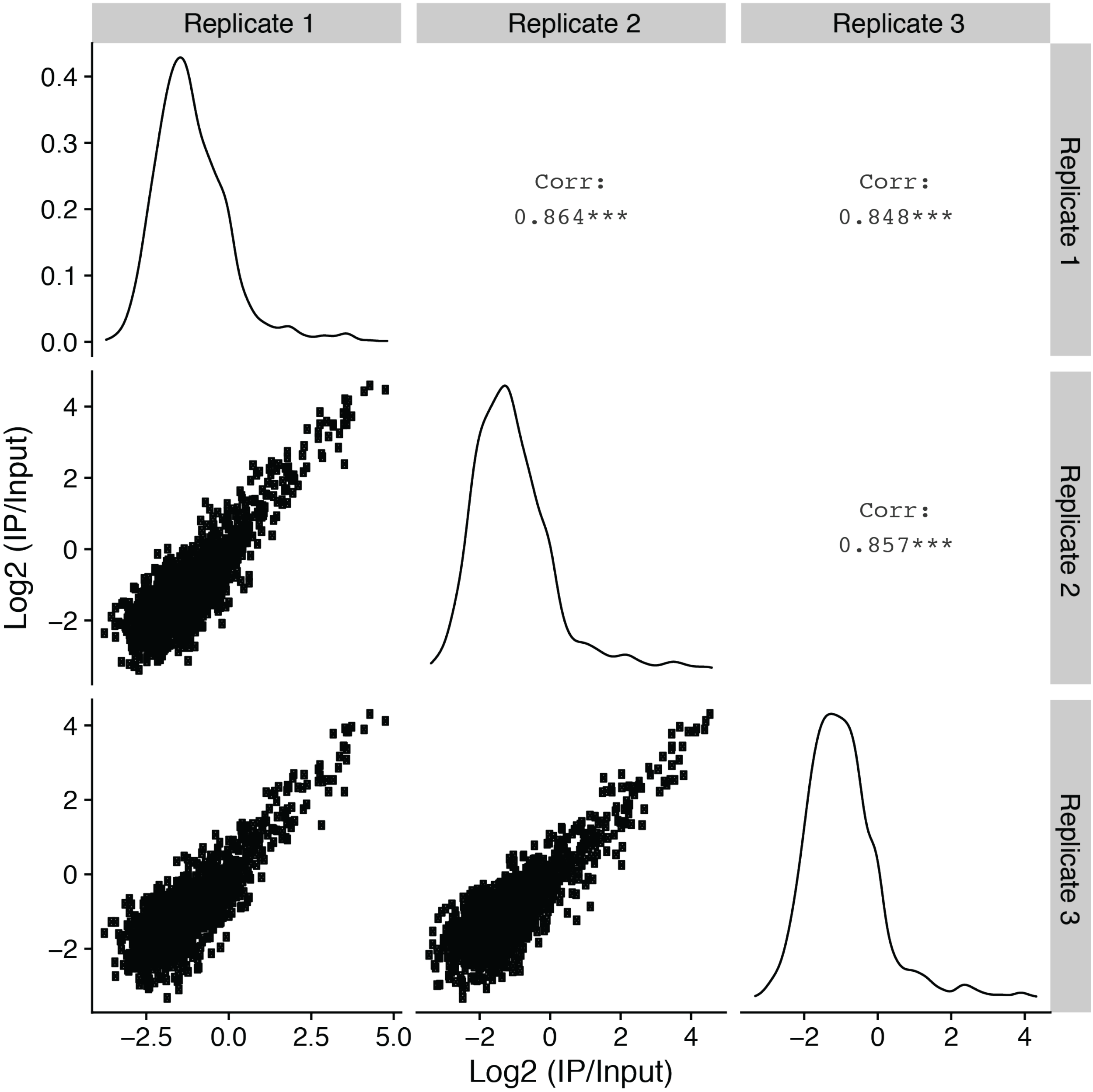


Enrichments of the tiles are highly correlated across three biological replicates. In the lower triangle, each dot represents a single tile. The distributions of log2 fold change (IP/Input) were shown in the diagonal. Correlation coefficients between two replicates were put in the upper triangle.

**Supplementary Figure 4. Volcano plot of MPRNA-IP for PUM2-binding RNAs**

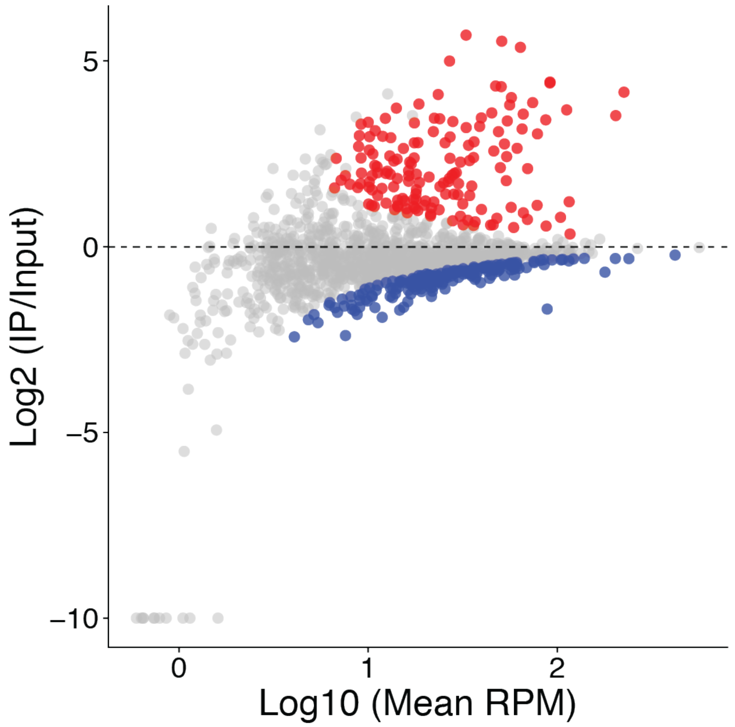


155 of the 1599 experimental sequences are significantly enriched (FDR adjusted p-value < 0.05 and IP/Input>1) in IP relative to Input. Red: enriched, blue: depleted, gray: insignificant.

**Supplementary Figure 5. Design of oligos in MPRNA-IP for MS2-MCP.**

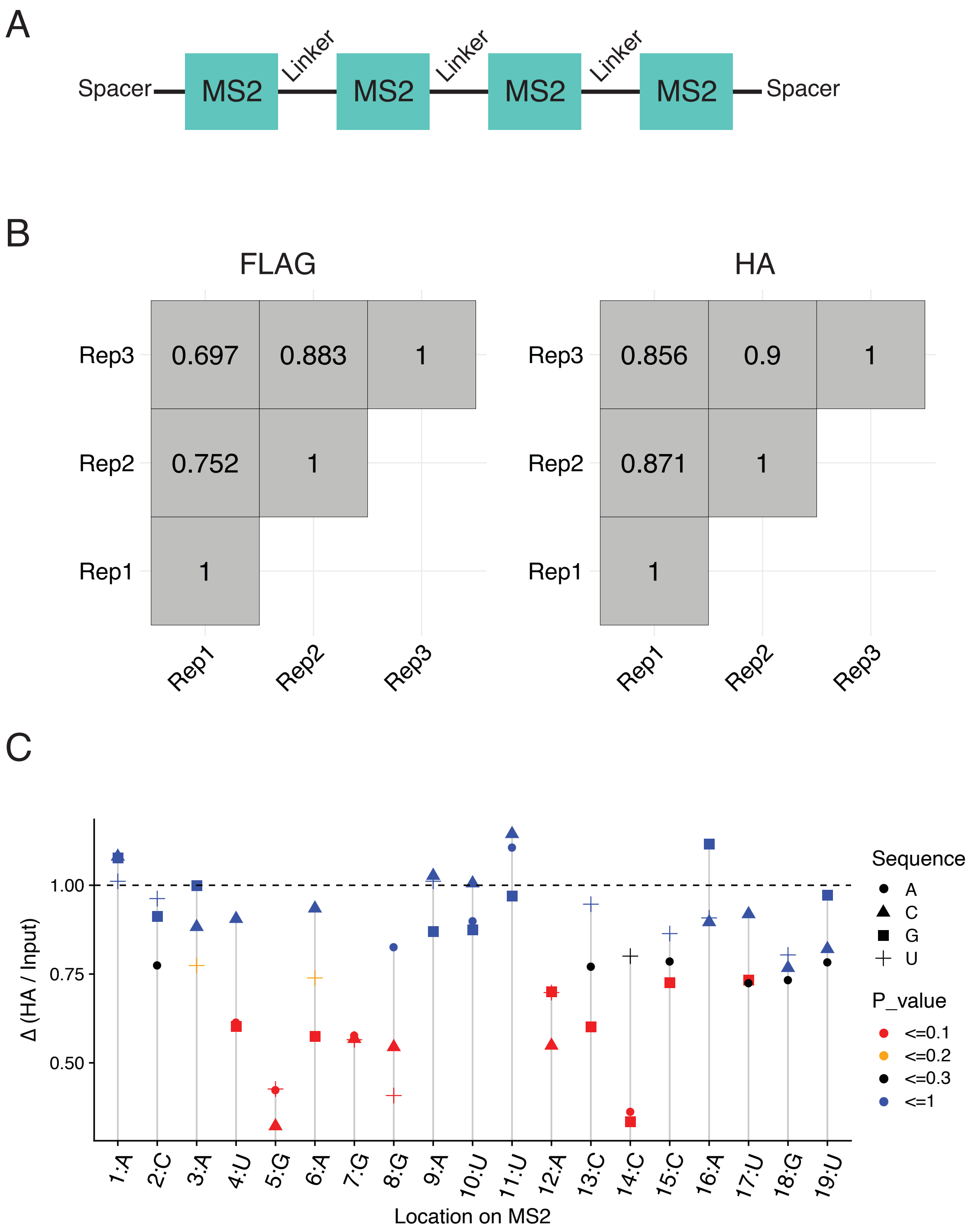


**(A)** In order to increase IP efficiency, each tile in this oligo pool included four copies of MS2 sequence, with unstructured linkers between neighboring hairpins. **(B)** Correlations across biological replicates in MS2-MCP MPRNA-IP: left for FLAG-MCP, right for HA-MCP. **(C)** The ratios of enrichment of the tiles with single mutation relative to WT (y-axis) are plotted along with MS2 WT sequence and location (x-axis). For a single location, three possible single mutations (denoted by shape) are described with their statistical significance (denoted by color).

**Supplementary Figure 6. MPRNA-IP for hTR-TERT.**

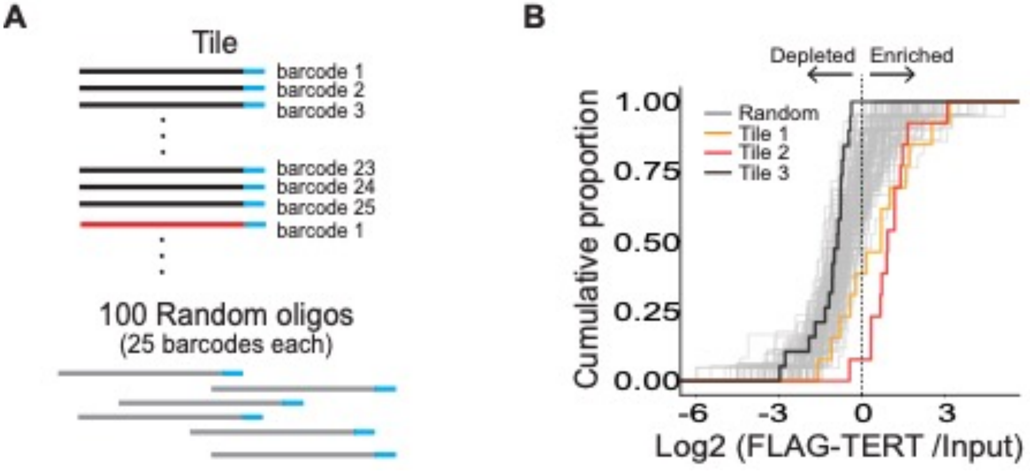


**(A)** Barcode multiplexing strategy for hTR pool design. Each tile is represented by 25 unique barcodes. Additionally, 100 random oligos were included as negative control sequences, each represented by 25 barcodes. This pool also contained mutants according to the compensatory mutational analysis strategy. **(B)** Tiles 1 and 2 are enriched by TERT IP, relative to both tile 3 and the 100 random sequences, confirming that the pseudoknot and CR4/5 domains are responsible for TERT interaction.

**Supplementary Figure 7. Compensatory mutation analysis using a four consecutive base-pair sliding window of mutation of hTR in MPRNA-IP of hTR-TERT.**

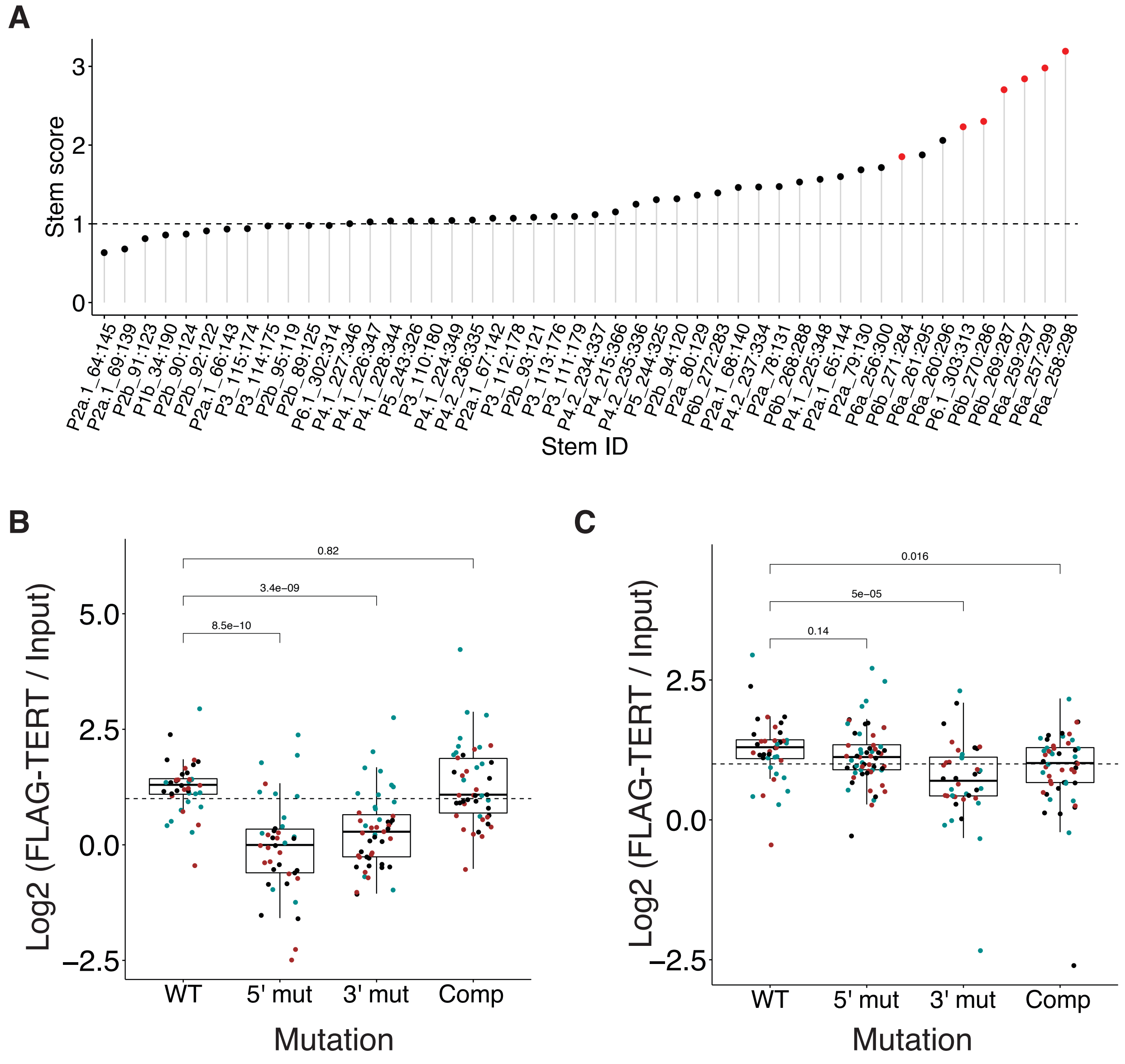


**(A)** Stem scores are described for 4-base-pair stems in pseudoknot and CR4/5 domains in hTR. Red is statistically significant stem while black is not (FDR < 0.05). **(B)** The enrichments of oligos (denoted by dots) of a stem in P6.1 (stem ID: P6.1_303:313) were described according to mutation types with three biological replicates (different colors). **(C)** Same as (B) for a stem in P5 (stem ID: P5_ 243:326).

**Supplementary Figure 8. Compensatory mutation analysis using single base-pair mutants of hTR in MPRNA-IP of hTR-TERT.**

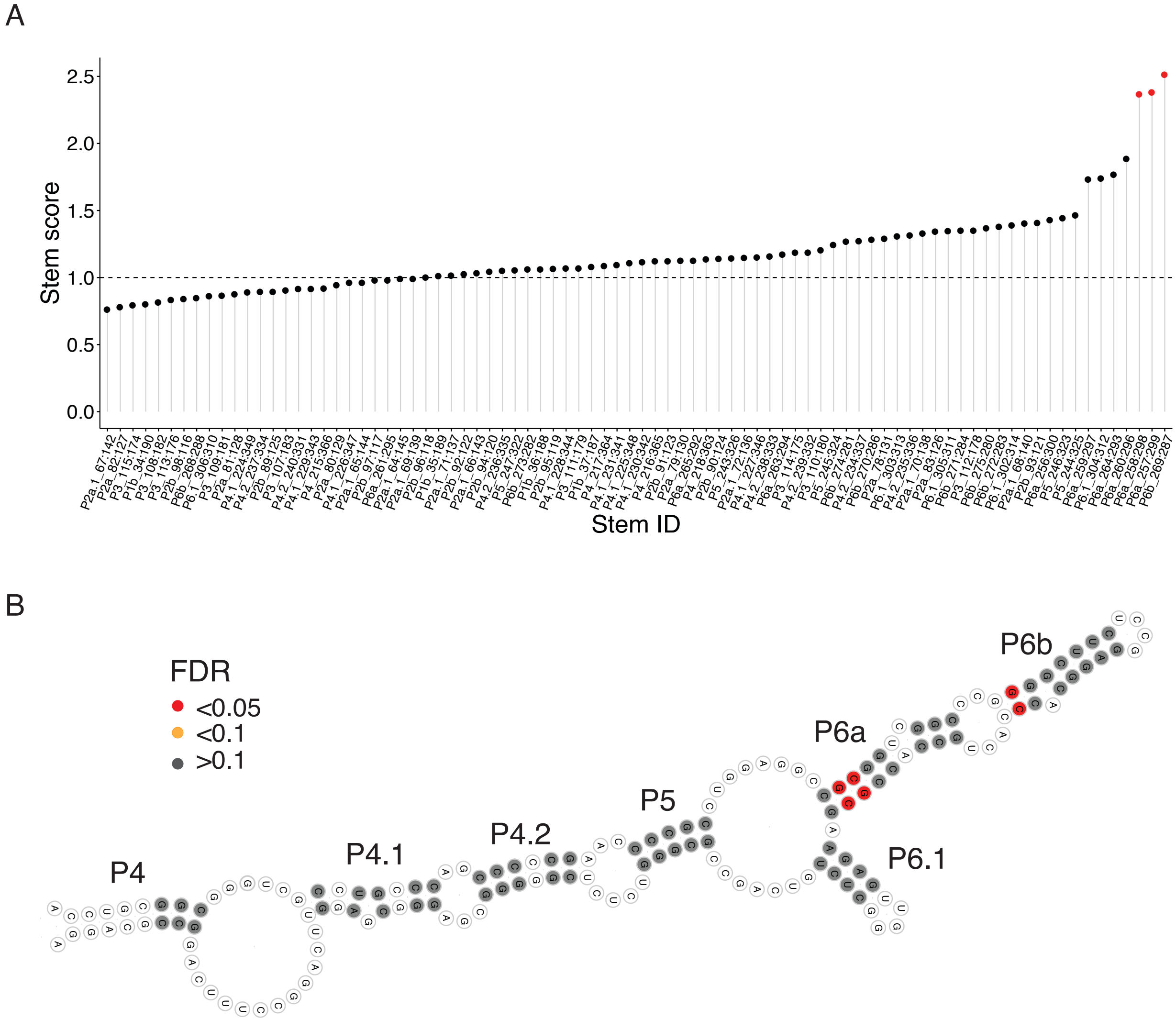


**(A)** Stem scores are described for single base-pair stems in pseudoknot and CR4/5 domains in hTR. Red is statistically significant stem while black is not (FDR < 0.05). **(B)** Statistical significances of stem scores of all known stems in tile 2 were overlaid on tile 2's secondary structure.

**Supplementary Table 1. Tiles with significant stem scores in hTR-TERT MPRNA-IP**

| No. | Mutation type | Region | Stem | 5' position | 3' position |
|---|---|---|---|---|---|
| 1 | 1bp | CR4/CR5 | P6a | 257 | 299 |
| 2 | 1bp | CR4/CR5 | P6a | 258 | 298 |
| 3 | 1bp | CR4/CR5 | P6b | 269 | 287 |
| 4 | 4bp | CR4/CR5 | P6.1 | 303 | 313 |
| 5 | 4bp | CR4/CR5 | P6a | 257 | 299 |
| 6 | 4bp | CR4/CR5 | P6a | 258 | 298 |
| 7 | 4bp | CR4/CR5 | P6a | 259 | 297 |
| 8 | 4bp | CR4/CR5 | P6b | 269 | 287 |
| 9 | 4bp | CR4/CR5 | P6b | 270 | 286 |
| 10 | 4bp | CR4/CR5 | P6b | 271 | 284 |
